# Acute and long-term circuit-level effects in the auditory cortex after sound trauma

**DOI:** 10.1101/2020.03.06.980730

**Authors:** Marcus Jeschke, Max F.K. Happel, Konstantin Tziridis, Patrick Krauss, Achim Schilling, Holger Schulze, Frank W. Ohl

**Author notes:** MJ and MFKH contributed equally to the study. HS and FWO contributed equally to the study. to whom correspondence should be addressed: Marcus Jeschke or Frank W. Ohl. Cognitive Hearing in Primates lab, Auditory Neuroscience and Optogenetics group, German Primate Center, Goettingen Germany; Institute for Auditory Neuroscience, University Medical Center Goettingen, Goettingen, Germany (Marcus Jeschke).

## Abstract

Harmful environmental sounds are a prevailing source for chronic hearing impairments, including noise induced hearing loss, hyperacusis, or tinnitus. How these symptoms are related to pathophysiological damage to the sensory receptor epithelia and its effects along the auditory pathway, such as functional reorganizations in the auditory cortex (ACx), have been documented in numerous studies. An open question concerns the temporal evolution of maladaptive changes after damage and their manifestation in the balance between afferent thalamocortical input and corticocortical input to the ACx.

To address this, we investigated the loci of plastic reorganizations across the tonotopic axis of the auditory cortex of male Mongolian gerbils (*Meriones unguiculatus*) acutely after a sound trauma and after several weeks. We used a laminar residual current-source density analysis to dissociate adaptations of intracolumnar input and horizontally relayed corticocortical input to synaptic populations across cortical layers in ACx. A pure tone-based sound trauma caused acute changes of subcortical inputs and corticocortical inputs at all tonotopic regions, particularly showing a broad elimination of tone-evoked inputs at tonotopic regions with a pre-trauma best frequency between 2-8 kHz. At other cortical sites, the overall columnar activity acutely decreased, while relative contributions of lateral corticocortical inputs increased. After 4-6 weeks, cortical activity to the altered sensory inputs showed a general increase of local thalamocortical input reaching levels higher than before the trauma. Hence, our results suggest a detailed mechanism for overcompensation of altered frequency input in the auditory cortex that relies on a changing balancing of thalamocortical and intracortical input and is confined to the spectral neighborhood of the trauma frequency.

**Significance statement:** Harmful noise exposure is a major anthropogenic cause of hearing disorders and is becoming an ever-increasing burden for human health and society. Damage to the sensory epithelia elicited by harmful sounds can subsequently lead to chronic hearing loss, hyperacusis or tinnitus. We still lack an understanding of the pathophysiological plastic processes and their evolution, particularly at the circuit level of the auditory cortex (ACx), which is fundamentally involved in auditory perception. We demonstrate that plastic changes in ACx after noise induced hearing loss (NIHL) occur over several weeks, and that changes in intracortical functional connectivity compensate the acute effects in the deafferentiated subcortical inputs. Such long-term changes may underlie the temporal evolution of hearing impairments or phantom sounds after NIHL.

## Introduction

Exposure to harmful environmental sound is a common cause for hearing impairments, including the development of noise induced hearing loss (NIHL) (Eggermont, 2017a, 2017b). Hearing loss has become increasingly prevalent over the last decades (Shargorodsky, 2010) and may further lead to hyperacusis, the increased loudness sensitivity to sounds, or tinnitus, the perception of phantom sound/s in the absence of physical sound sources. Exposure to high sound levels can severely damage the peripheral sensory receptor epithelia within the cochlea which presents the most reasonable initial factor for such pathological disturbances: Cochlear damage causes altered auditory processing and, in consequence, may lead to compensatory and/or erroneous functional map plasticity in higher sensory brain areas, as for instance the auditory cortex. Such maladaptive map plasticity has been proposed to underlie the development of central tinnitus (Eggermont, 2003; Engineer et al., 2011), although these classical models of tinnitus development have recently been challenged by alternative models (Schaette and Kempter, 2006; Krauss et al., 2016). In humans, as well as in animal models, plastic changes of the tonotopic organization of the auditory cortex (ACx) have been found after noise exposure (Muhlnickel et al., 1998; Eggermont and Komiya, 2000; Dietrich et al., 2001; Norena, 2005; Chen et al., 2016). However, until now cortical map plasticity has been mainly documented by comparing tonotopic organization before and after noise exposure (Eggermont, 2017b). Therefore, we still do not fully understand the temporal evolution of plastic processes within the cortical network following noise trauma. On the other hand, a deeper understanding of such circuit-level effects might be important to understand the temporal and/or permanent perceptual threshold shifts induced by environmental sounds and their relation to transient and chronic forms of hyperacusis, hearing loss and tinnitus.

To achieve such understanding of the maladaptive plastic changes after noise exposure along the auditory pathway, it would be important to describe the loci of plastic reorganizations from peripheral to subcortical and cortical regions of sensory processing. Whereas peripheral damage initially leads to reduced activity within the damaged region of the cochlea, increased activity has been described for several nuclei along the auditory pathway from the dorsal cochlear nucleus on (Kaltenbach et al., 1998, 2004; Kaltenbach and Afman, 2000; Brozoski et al., 2002; Zacharek et al., 2002; Sun et al., 2009; Holt et al., 2010; Wu et al., 2016) and was interpreted as a potential physiological correlate of tinnitus (e.g. Noreña and Eggermont, 2003; Engineer et al., 2011; Ahlf et al., 2012; Tziridis et al., 2015). These findings seem to suggest that reduced input within the peripherally deafferented frequency range is not only compensated along the auditory pathway, but eventually leads to hyperactivity in specific tonotopic regions within the auditory pathway. It has been hypothesized that this phenomenon might be explained by selective amplification of previously subthreshold cortical inputs which then provide the major contribution to cortical activation during tinnitus (Irvine, 2018). Other models explain the subcortical enhancement of activations by means of homeostatic plasticity (Schaette and Kempter, 2006; Schaette and McAlpine, 2011) or stochastic resonance (Krauss et al., 2016). Gathering experimental data with respect to the involved specific local and corticocortical synaptic populations in cortex during such plastic adaptations to altered cochlear input is technically challenging and hence still elusive. In a previous study we developed a method to disentangle activations of neuronal populations from thalamocortical or intracortical sources using a relative residual analysis of the current-source density (CSD) distribution (Happel et al., 2010). Of particular interest for the description of the development of maladaptive adaptations to NIHL such as tinnitus is the time point of observation after noise trauma, which has not been quantitatively analyzed so far.

Therefore, the goal of this study was to utilize the analysis of the relative residual of the CSD and to disentangle the short- and long-term plastic effects of extensive sound exposure on thalamic input, local cortical processing and long-range corticocortical networks at the level of the primary auditory cortex (AI). Thereby, we aimed at a better understanding of the circuit effects following exposure to loud sounds and to investigate potential compensatory mechanisms and/or chronic maladaptive processes that for instance may be responsible for the manifestation of a central tinnitus. Our data suggest that a pure tone-based sound trauma causes acute changes of subcortical inputs and corticocortical circuits at all tonotopic regions in auditory cortex. Long-lasting adaptive processes adjust cortical processing over weeks to the altered sensory inputs. A better understanding of the circuit mechanisms underlying this dynamic temporal process may provide new implications to ameliorate the effects of noise-induced hearing loss or its common perceptual symptom of chronic tinnitus.

## Materials and Methods

Experiments were performed on 17 adult male Mongolian gerbils (*Meriones unguiculatus*) (age: 3 – 6 months, body weight: 70 – 120 g) anesthetized with ketamine-xylazine. All experimental procedures were approved by local authorities of the State of Saxony-Anhalt, Germany, and were in accordance with the international guidelines for care and use of animals in research as detailed by the NIH.

### Surgical procedure

Anesthesia was induced by intraperitoneal infusion (0.06 ml/h) of 45 % ketamine (50 mg/ml, Ratiopharm, Germany), 5 % xylazine (Rompun, 2 %, BayerVital, Germany) and 50 % isotonic sodium chloride solution (154 mmol/l, Braun, Germany). Throughout the anesthetic procedures the animal’s body temperature was maintained at 37 °C by means of remote controlled heating pads. Anesthetic depth was tested periodically using the hind limb withdrawal reflex. To reduce tracheobronchial secretions Glycopyrrolate (Robinul, 0.02 ml SC) was used after surgery and prior to recording. To fixate the animals head during auditory free field stimulation we glued two M3 standoffs onto the midline of the skull using light curing adhesive and composite (Plurabond One-SE and Plurafill Flow, Pluradent, Offenbach, Germany) as a base layer and dental acrylic to shape the skull implant. Next, access to the right (n=2) or –bilateral (n=15) ACx was achieved by removing the temporal muscle, followed by a craniotomy overlying the ACx between the eye and ear. A stainless steel insect pin placed in the frontal bone with contact to the dura served as reference for electrophysiological recordings.

### Electrophysiological recordings

After surgery, animals were transferred into an electrically and acoustically shielded sound-proof chamber (IAC, Niederkrüchten, Germany). The animals head was fixed in place facing a speaker (Tannoy arena satellite; distance: 1 m) by screwing the head cap to a custom-made head holder. Using vasculature landmarks (Thomas et al., 1993) laminar 32 channel silicon arrays (Type: A1×32-5mm-50-413, Neuronexus Technologies, USA) were slowly inserted orthogonally to the cortical surface into field AI of the ACx through small slits in the dura made by hypodermic needles. After insertion, the tissue was allowed to settle for at least 20 min. Local field potential (LFP) responses to pure tones covering a frequency range from 0.5 to 32 kHz (half-octave or octave steps; 100 ms duration; at least 80 repetitions) at a sound level of 44 and 64 dB SPL were recorded (MAP system, Plexon Inc, USA) to determine the best frequency (BF) at the recording site. Next, responses to varying sound levels (covering −6 to 94 dB SPL in 10 dB steps) at several frequencies were recorded. These frequencies included the BF, the sound trauma frequency (2 kHz, see below), one frequency at least 1 octave below and above the trauma frequency. All sounds were generated in Matlab, converted to analog signals with a data acquisition card (NI PCI-BNC2110; National Instruments, Germany), fed through a programmable attenuator (g.PAH, Guger Technologies; Austria) and amplified by a wide-range audio amplifier (Thomas Tech Amp75). A measurement microphone and conditioning amplifier were used to calibrate acoustic stimuli (G.R.A.S. 26AM and B&K Nexus 2690-A, Bruel & Kjaer, Germany).

### Sound trauma and experimental time line

For sound trauma induction a sine wave generator (HP 33120A) generated a 2 kHz sine wave which was amplified by a power amplifier (Alesis RA150) and fed to a separate speaker (Canton XS.2) placed 20 cm in front of the animal. The stimulation lasted for 75 min. Sound level was calibrated for each trauma induction to a final level of 115 dB SPL as explained above. Such trauma has been shown to produce a mild to moderate hearing loss in the range of 20 to 40 dB around the trauma frequency (Ahlf et al., 2012). Sound evoked responses from field AI of the ACx were recorded immediately before and after sound trauma induction (**Fig. 1A**) within the same recording session. Afterwards, craniotomies were filled with antibiotic gel (K-Y jelly) and covered by dental cement. Animals were then allowed to recover and underwent a second, final, recording session 4-6 weeks later. Again, animals were anesthetized, the craniotomies reopened and animals were transferred to a sound proof chamber. Location of recording site during trauma induction and during the recovery measurement 4-6 weeks later was kept similar due to the vascularization pattern, which serve as reliable landmark for the tonotopic gradient in gerbil ACx (Thomas et al., 1993; Schulze et al., 1997).

**Fig 1:**
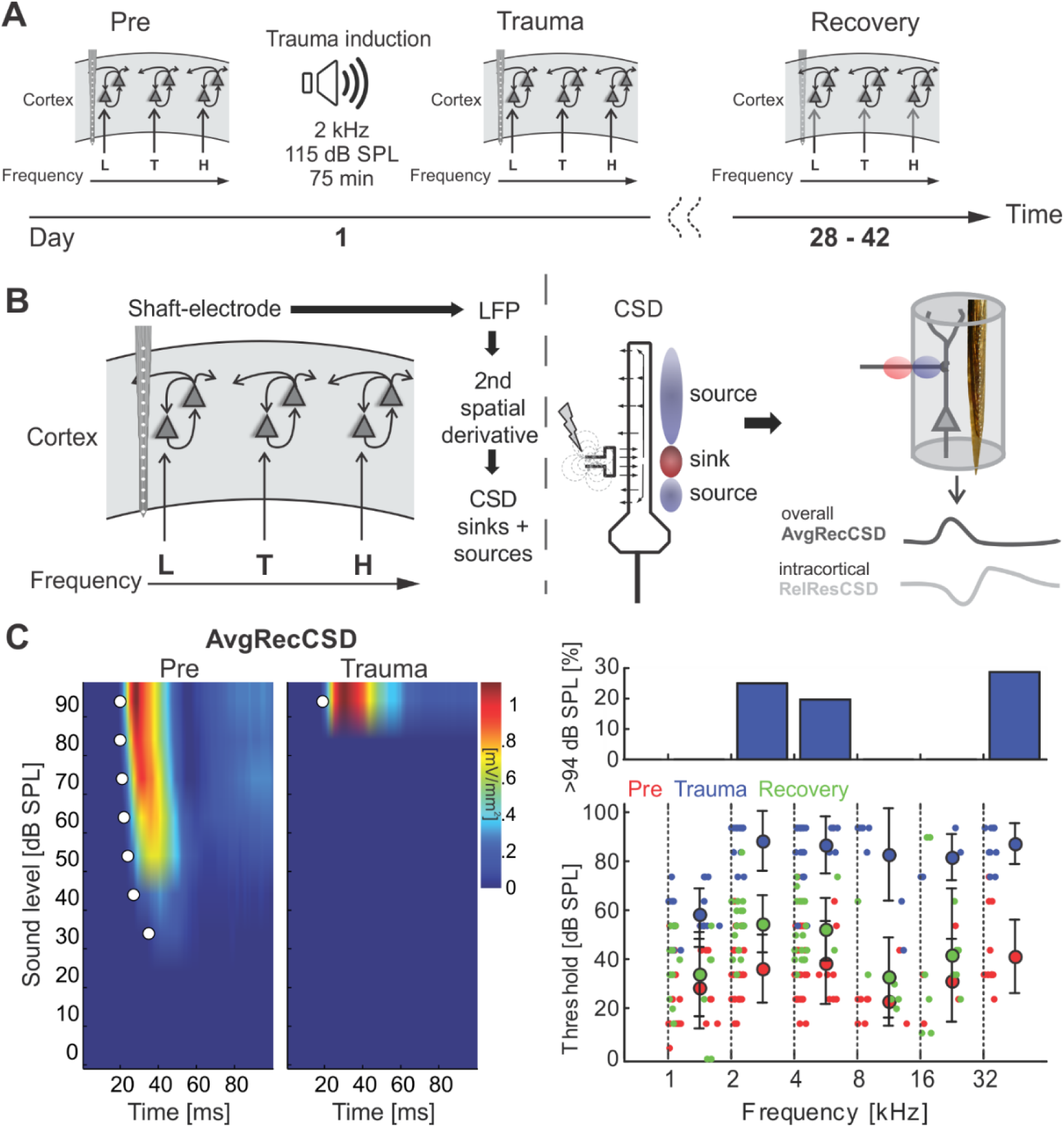
**A)** Schematic representation of the timeline of experiments investigating sound trauma related effects in the auditory cortex. Electrophysiological recordings were obtained directly before (*Pre*) and after trauma (*Trauma*) as well as after 4 to 6 weeks (*Recovery*). The schematics symbolize a piece of auditory cortex with an implanted shaft electrode and tonotopic locations corresponding to the trauma frequency (T) as well as lower (L) and higher frequencies (H). **B)** Schematic of data analysis strategy (see main textfor further explanation). **C)** Analysis of significant AvgRecCSD responses to investigate sound trauma related effects on cortical response thresholds. A representative example (left panel) depicts the AvgRecCSD (heat map coded; warm colors correspond to larger AvgRecCSD values; significant responses are indicated by white circles) in response to presentations of 2 kHz tones and illustrates the increase in threshold from 34 dB SPL to 94 dB SPL following trauma induction. Thresholds determined in the population data were plotted for different frequency bins and time points relative to trauma (right bottom panel). Immediately after trauma induction (blue circles) a threshold could not be determined in a number of cases (right top panel). Four to 6 weeks after trauma thresholds (green circles) recovered to a large degree.

### Data analysis

Based on the second spatial derivative of the laminar LFP profiles recorded from the auditory cortex in response to pure tone stimulation (**Fig. 1B**) we calculated one-dimensional current source density (CSD) distribution as:

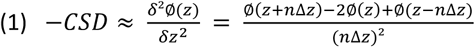

where Φ is the field potential, z the spatial coordinate perpendicular to the cortical laminae, Δz the spatial sampling interval (50 µm) (Mitzdorf, 1985). LFP profiles were smoothed with a weighted average (Hamming window) of 5 channels (corresponding to a spatial filter kernel of 250 µm; linear extrapolation of 2 channels at boundaries; see (Happel et al., 2010). CSD profiles reveal patterns of current influx (sinks) and efflux (sources). From single trial CSD profiles we computed the average rectified CSD (AvgRecCSD) to investigate the temporal pattern of the overall transmembrane current flow at the recorded site (Givre et al., 1994) as:

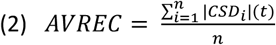

The relative residue of the CSD (RelResCSD) is defined as the sum of the non-rectified magnitudes divided by the rectified magnitudes for each channel:

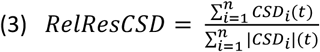

Thereby, the RelResCSD quantifies the balance of the transmembrane charge transfer along the recording axis (Harding, 1992) and gives rise to the lateral corticocortical contribution to stimulus related activity (Happel et al., 2010).

The tonotopic region of the cortical patch under observation was characterized based on responses to varying pure tones covering a frequency range from 0.5 to 32 kHz at moderate sound levels (44 and 64 or 64 dB SPL). Based on the canonical CSD pattern we defined individual sink components in granular (S1), supragranular (S2) and early (iS1) and late (S3) infragranular layers, as explained in detail elsewhere (Happel et al., 2010); for an example, see Fig. 5A, top left). Peak amplitudes and onset latencies of individual current sinks were determined for individual channels and then averaged. Onset latencies were determined using a linear fit around the point where each curve surpasses 3 standard deviations above/below baseline (Happel et al., 2010). Frequencies evoking maximal responses of the granular sink were defined as the BF. Response threshold was determined as the lowest sound intensity eliciting a significant response at any characteristic frequency 3 standard deviations over baseline (> 5ms). Response bandwidths were quantified as Q40dB-values 40 dB above response threshold.

**Fig 2:**
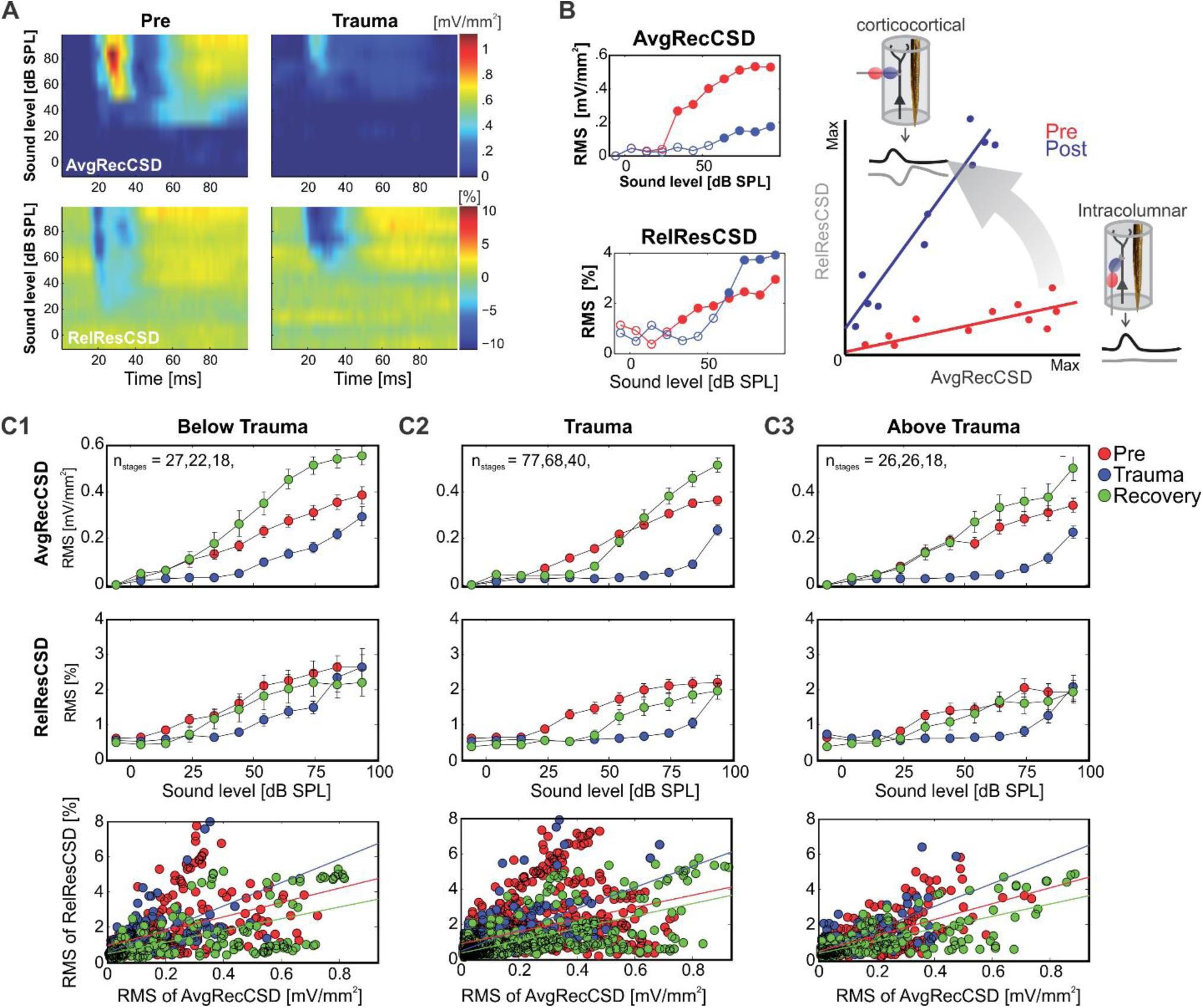
**A)** Representative example of AvgRecCSD (top panels) and RelResCSD (bottom panels) evoked by 1 kHz stimulation prior to (left) and after trauma induction (right) at a cortical site with a BF of 5.6 kHz. **B)** For further quantitative analysis root-mean-square values during 100 ms stimulus presentation were calculated for AvgRecCSD and RelResCSD. Prior to trauma induction (red circles) the threshold of activation was 34 dB SPL (filled circles) and increased to 64 dB SPL after trauma. While AvgRecCSD values generally decreased after trauma, RelResCSD values even increased from around 2 to 4 %. Note, that both before and after trauma induction AvgRecCSD and RelResCSD were highly correlated (p < 2*10^−5^). In the schematic depiction of the comparison between both parameters (*right*), red and blue lines indicate linear regression lines before and after trauma, respectively. Comparison of their slopes and offsets allow us to investigate the effect of sound-trauma on local and corticocortical synaptic circuits. **C)** Across all experiments and independent of the frequency of pure tone stimulation relative to the trauma frequency (**C1-3**) AvgRecCSD (top panels) values increased with increasing sound levels (data are shown as mean ± SEM). As for the individual example in A) and B) RelResCSD (middle panels) values also increased with increasing level. Expectedly, after trauma a reduction of AvgRecCSD values was observed. Interestingly, several weeks after trauma (green circles – Recovery), higher mean AvgRecCSD values than before trauma were observed. These changes seem to be not counterbalanced by relative increases in RelResCSD values. A regression analysis on the relationship between AvgRecCSD and RelResCSD revealed significant correlations in all cases analyzed (p < 10^−13^). For further quantitative analysis see main text.

**Fig 3:**
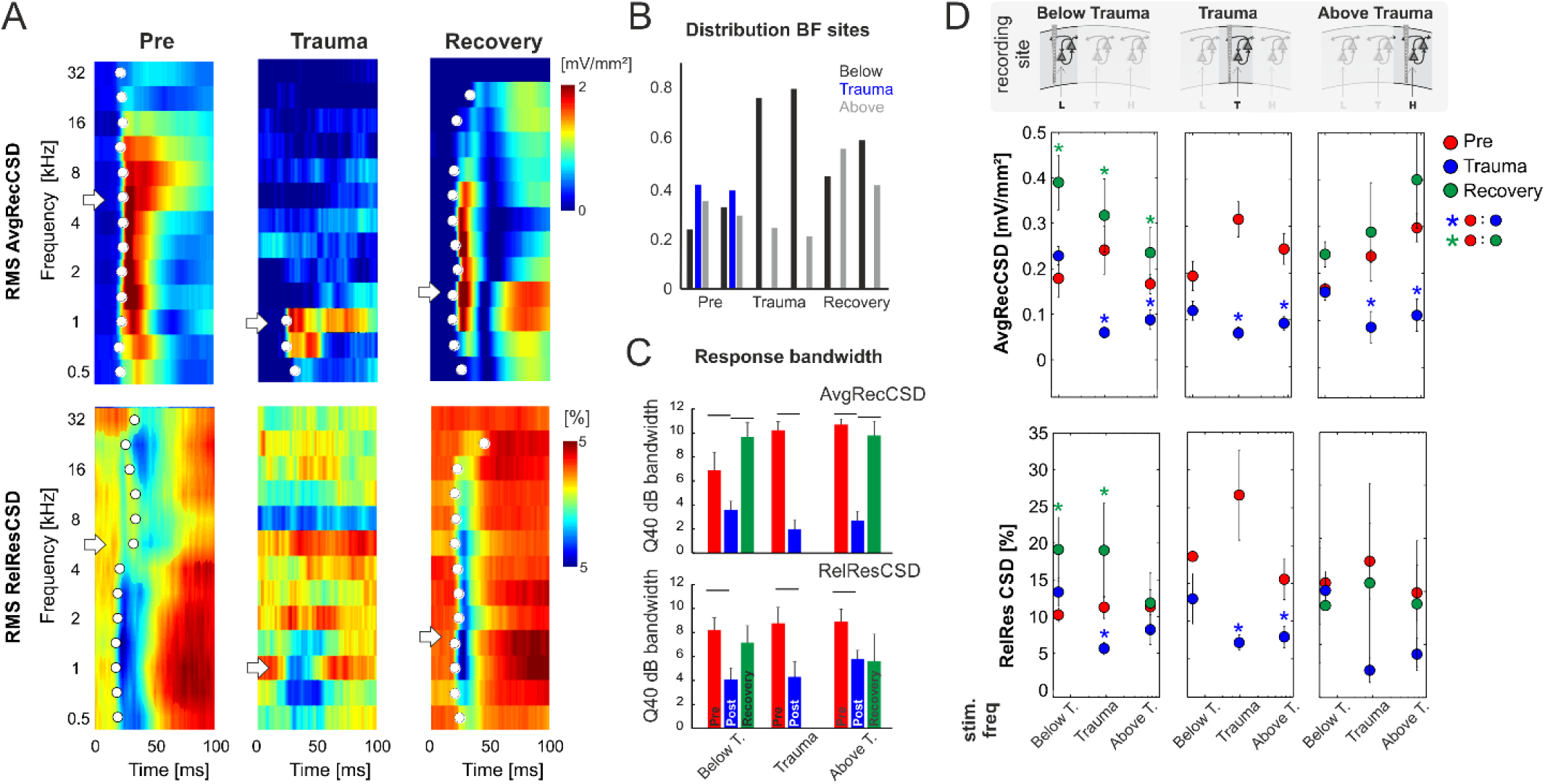
**A)** Representative example of AvgRecCSD (top) and RelResCSD (bottom) obtained after stimulation with varying sound frequencies at moderate sound level (64 dB SPL) prior to (left), after trauma induction (middle) and after 4 weeks of recovery (right). The respective BFs defined by the maximum AvgRecCSD peak amplitude are indicated by white arrows. Before trauma, the BF in the example was found to be 5.6 kHz. At the corresponding recording position, the BF after 4 weeks of recovery was decreased to 1.4 kHz. Onset latency of significant activation (2SD > baseline) at each sound frequency is indicated by white circle. **B)** Distribution of the BF (based on granular sink S1 peak amplitude) measured at recording sites with BFs below, around and above the trauma frequency (bar colors) at the three time points directly before and after trauma induction as well as after 4 weeks of recovery. For ChiSquare-test results, see main text. **C)** Response bandwidths before (*Pre*) and after (*Trauma*) the trauma and 4 weeks later (*Recovery*) measured by Q40dB values (in octaves) were calculated for the RMS values of the AvgRecCSD (top) and the RelResCSD (bottom). **D)** Quantitative comparison of RMS values of AvgRecCSD (top) and RelResCSD (bottom) obtained after stimulation at moderate sound level (64 dB SPL) with sound frequencies below, at and above the trauma. Data prior to (red), after trauma induction (blue) and after 4-7 weeks of recovery (green) are plotted. For statistical results and further explanation see main text. Blue and green asterisks mark significant differences between groups identified by post-hoc paired-sample Student’s t-test following rmANOVA analysis (see **Tab. 3.1-3.5** based on a Bonferroni-corrected level of significance due to multiple testing).

**Fig 4:**
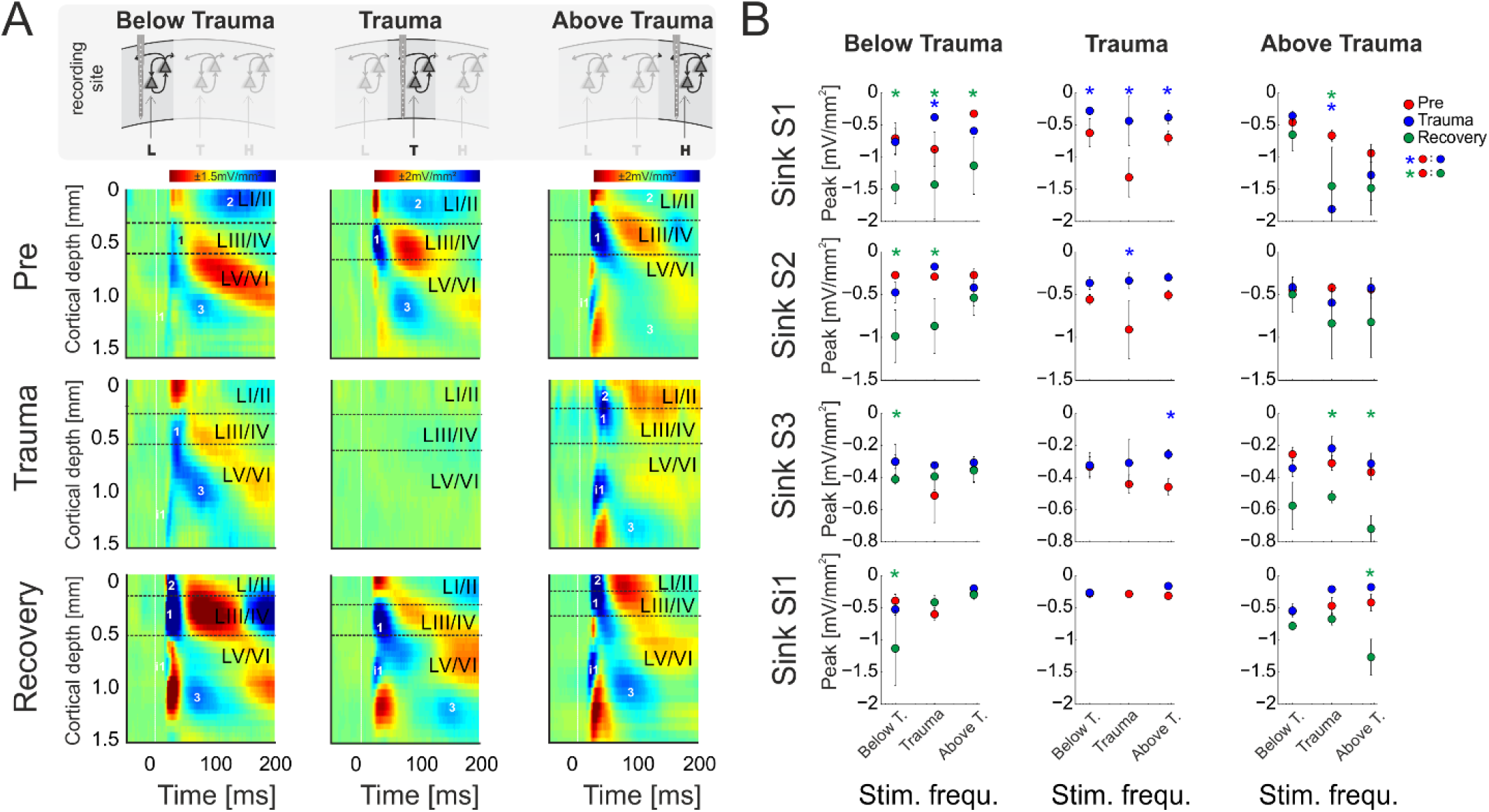
**A)** CSD profiles obtained at a moderate sound level (64 dB SPL) before (*Pre*) and after trauma induction (*Trauma*) as well as after 4 - 6 weeks of recovery (*Recovery*) for all three categories of recording sites, viz. with BF representations below, around and above the trauma frequency (left to right). Effects of sound trauma induction on respective sink components S1, S2, S3 and iS1 are shown for stimulation with the BF from pre-measurement. **B)** Quantification of sink peak amplitudes prior to (red), after trauma induction (blue) and after 4 weeks of recovery (green) at the different recording sites and after stimulation with varying sound frequencies (Low, Middle, High). For statistical results and further explanation the reader is referred to the main text. Blue and green asterisks mark significant differences between groups identified by post-hoc paired-sample Student’s t-test following rmANOVA analysis (see **Tab. 3.6-3.8** based on a Bonferroni-corrected level of significance due to multiple testing).

**Fig. 5:**
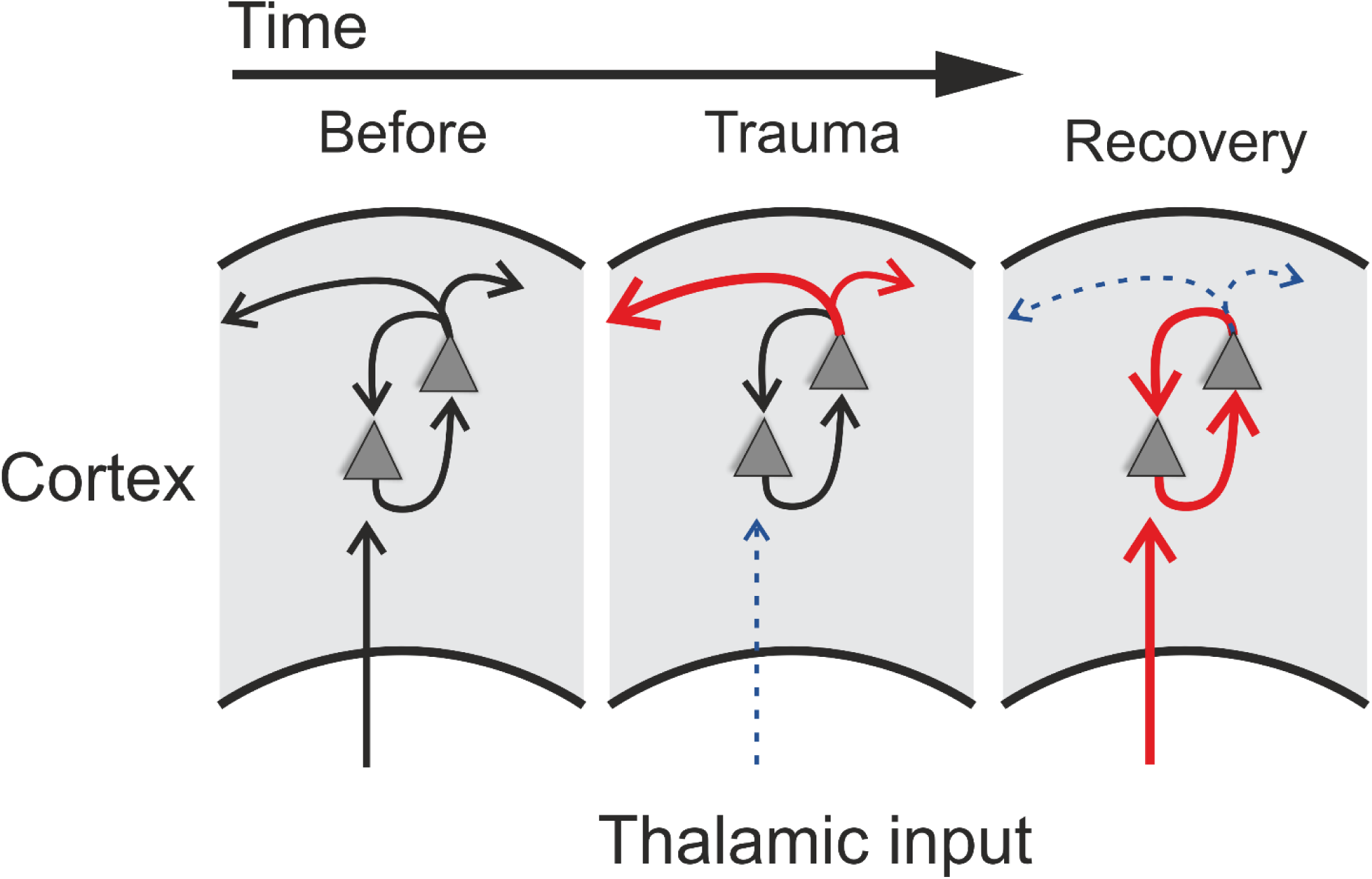
Schematic illustration of trauma-induced changes over time. Note that lateral connections in the schemes are symmetric to higher and lower frequency regions, but for clarity we sketched only one direction. Acoustic trauma led to increased auditory thresholds over the entire tonotopic gradient (cf. **Fig. 1C**). On a columnar level, this can be explained by decreased strength of local thalamic input to ACx (indicated by dashed thin blue dashed arrow in middle panel; cf. **Fig. 3A,D**). While overall columnar activity was consistently decreased across the tonotopic gradient, the relative contribution of corticocortical activity was increased immediately after the trauma (red arrow in middle panel; cf. **Fig. 2**). After recovery over weeks, these acute effects were reversed: while increased sensory input from thalamus was now coupled with an increased local gain of tone-evoked cortical activity (red arrows in right panel; cf. **Fig. 4**), the recruitment of lateral corticocortical circuits was relatively decreased to levels even below (for frequency regions above trauma, blue dashed arrow in right panel) or equal to pre-trauma conditions (cf. **Fig. 2**).

For further quantitative analyses, we split the tonotopic axis into tonotopic regions below the trauma (0.5 – <2 kHz), around the trauma (2 – <8 kHz) and above the trauma (8 – 32 kHz). Thresholds for pure tone stimulation were calculated based on sound intensity profiles of AvgRecCSD responses and were taken as the lowest sound level at which the root-mean-squared AvgRecCSD during stimulus presentation exceeded 3 standard deviations above prestimulus baseline levels. RelResCSD were considered if the AvgRecCSD response was found to be significant. The relationship between the overall strength of cortical activation and potential corticocortical contributions was investigated by correlating the AvgRecCSD and RelResCSD obtained from sound intensity profiles. The slope and offset of linear regression lines were then taken as indicators of corticocortical contribution to cortical activity at a given recording site. Specifically, a steeper slope hence indicates a higher corticocortical contribution with increasing overall activation. The offset, however, reveals the strength of corticocortical contribution at the minimal cortical activation.

### Statistical analysis

To investigate potential associations between two variables a Pearson correlation analysis was performed. In order to further analyze the relationship between two variables and compare potential changes across experimental conditions linear regressions were calculated. Parameters of linear regressions were compared with general linear models with the factors experimental stage (*Pre, Trauma, Recovery*) and tonotopic region (*below trauma, trauma, above trauma*). For statistical comparison between two groups, we used paired-sample Student-t-tests. Comparison of multiple groups was performed by multi-factorial repeated measures ANOVAs (with Huynh-Feldt correction of sphericity) with a general significance level of α = 0.05. For post-hoc tests we used paired-sample Student-t-tests. with Bonferroni-corrected levels of significance level in case of testing repeatedly for n_test_ times of α* = α / n_test_.

## Results

### Acoustic trauma led to increases in threshold of cortical activation

We here investigated the effect of a permanent mild to moderate sound-induced hearing loss on auditory cortical processing. To induce the acoustic trauma we employed a previously established procedure, presenting a 2 kHz pure tone at 115 dB SPL for 75 min (Ahlf et al., 2012). Using implanted multichannel shaft electrode arrays, penetrating all cortical layers, electrophysiological recordings of local field potentials evoked by pure tones of varying frequency and intensity were performed immediately before and after trauma induction as well as after 4-6 weeks of recovery (**Fig. 1A**). From the local field potential data, current source density (CSD) profiles across the cortical laminae were reconstructed, and in addition averaged rectified CSDs (AvgRecCSD) and relative residuals (RelResCSD) were determined as measure of the overall cortical activity in a cortical column around the shaft electrode and estimate of the relative contribution of intracortical horizontal input to this activation, respectively (see (Happel et al., 2010) (**Fig. 1B**). Analysis of the cortical activation using pure-tone evoked responses (**Fig. 1C**, **left**) revealed that at the trauma frequency thresholds prior to trauma induction were 36.4 ± 13.7 dB SPL and increased to 88.2 ± 12.1 dB SPL acutely after trauma resulting in an average increase in threshold of 51.8 dB (**Fig. 1C**). However, this only serves as a lower estimate as we were not able to drive significant cortical activation at the highest sound level tested (94 dB SPL) in 25 % of recordings (**Fig. 1C, upper right panel**). Threshold increases were found throughout a large frequency region extending from at least 1 octave below and up to 4 octaves above the trauma frequency. At 1 kHz thresholds increased by an average of 30.1 dB from 28.4 ± 16.5 dB. From 4 kHz to 32 kHz thresholds increased on average by 50.9 ± 6.3 dB. Even at a stimulation frequency of 32 kHz we were not able to determine a response threshold immediately after the trauma in 4 out of 14 cases. When tested after several weeks, thresholds recovered to a large extent but not completely. A two-way repeated measures ANOVA with factors “experimental time point” and “frequency bin” yielded a main effect for the “time point” (*Pre* vs. *Recovery*; F_1,195_ = 19.277, p < 1.85*10^−5^) such that average thresholds were 45.13 ± 18.6 dB after recovery amounting to an average residual threshold increase of 10.1 dB. Additionally, the main effect of the factor “frequency bin” was also significant (F_5,195_ = 6.729, p < 8.268*10^−6^).

### Acute increase of corticocortical activity after sound trauma and long-term compensation

The relative weights of thalamocortically and intracortically relayed contributions, respectively, to cortical plasticity following exposure to intense sounds are yet unclear.. The analysis of the relative residues of the CSD allows testing whether the recruitment of corticocortical horizontal processes is altered following acoustic trauma. We followed the main hypothesis that after sound trauma the RelResCSD (which is mainly determined by intracortically, rather than thalamocortically, relayed inputs) is increased even though the overall cortical activation might be diminished. **Fig. 2A** and **B** illustrate a representative example in which the overall cortical activation was decreased after trauma induction (1 kHz stimulation; BF of 5.6 kHz) while, in line with this hypothesis, the contribution of corticocortical horizontal input was increased. This can be further assessed by calculating the slope of the regression between the AvgRecCSD and the RelResCSDamplitudes which indicates how strongly corticocortical contributions increase with increasing overall activation (**Fig. 2B; right**). We calculated the root mean square values during stimulus presentation for AvgRecCSD and RelResCSD to quantify our observations. Although the threshold of activation increased by 30 dB with an accompanying decrease of the strength of activation by 77 %, the relative corticocortical contribution increased by ∼10 %. Above threshold this pattern remained: while the strength of activation was decreased (by 71 – 67 % from 64 to 94 dB SPL) the relative corticocortical contribution consistently increased (by 51 – 33 % from 64 to 94 dB SPL). Notably, as the strength of activation increased with increasing sound level, so did the corticocortical contribution. Consequently, the AvgRecCSD and RelResCSD were highly and significantly correlated before (R = 0.943; p < 1.4*10^−5^) and after trauma induction (R = 0.95; p < 9*10^−6^). Across all experiments (**Fig. 2C**) and independent of the frequency of pure tone stimulation the overall strength of cortical activation (**Fig. 2C; top**) increased with increasing sound level in all experimental phases (Pre, Trauma and Recovery). After trauma induction a reduction of cortical activation was observed consistent with earlier observations of a reduced subcortical drive (Heeringa and van Dijk, 2016). Furthermore, the relative corticocortical contribution also increased with increasing level (**Fig. 2C; middle**). However, several weeks after the trauma, stronger cortical activation than before trauma was observed, while the intracortical contribution did reach values as approximately before the trauma (although there were slight differences above and below trauma frequency regions, see below). Therefore, we conclude that plastic reorganizations over weeks seem to have compensated for the acute increased lateral, corticocortical spread of activity.

Since significant correlations (p < 10^−13^; **Fig. 2C bottom**) between the strength of cortical activation and corticocortical contributions were observed during all phases of the experiment, we did not directly compare the data before and after trauma induction. Instead, we performed a linear regression analysis between AvgRecCSD and RelResCSD values to reveal potential changes in the recruitment of corticocortical processes after sound trauma independent of the strength of cortical activation (**Fig. 2C**). The slope of the linear regression between both measures, as indicator of the corticocortical contribution (see **Fig. 2B**), was steeper immediately after trauma (blue), while similar slopes were observed between pre-trauma (red) and recovery (green; detailed statistical analysis in the next paragraph). In other words, immediately after trauma induction, corticocortically relayed synaptic activity contributed more to the overall activation strengths. This relative increase recovered over several weeks (**Fig. 2C bottom**). In order to quantify these observations we fitted linear regression models to the AvgRecCSD and RelResCSD data to obtain measures of slope and offset of these regressions. The linear regression analysis indicated that the slope between AvgRecCSD and RelResCSD, i.e. the contribution of corticocortical activity to the overall cortical activity, is dependent on the experimental stage and tonotopic region (R^²^ = .08, adjusted R^²^ = 0.07, F_4,297_ = 6.491, p < 5.1*10^−5^). Profound inter-subject differences probably explain the remaining variability. A trend for increased slopes was observed immediately after trauma (ß = 1.31, p = 0.059), whereas lower slopes were detected after several weeks of recovery (ß = −2.04, p = 0.008). A trend was further found for increased slopes in the trauma region (ß = 1.55, p = 0.054). For further details see **Tab. 1**. The offset between AvgRecCSD and RelResCSD also depended on experimental stage and tonotopic region (R^²^ = .03, adjusted R^²^ = 0.02, F_4,297_ = 2.578, p = 0.038). The remaining variability is potentially explained by large differences between subjects. A trend for decreased offsets was observed immediately after trauma (ß = −0.11, p = 0.096) as well as increased offsets in the region above the trauma (ß = 0.17, p = 0.008). For further details see **Tab. 2**.

**Table 1.** Linear regression of slope. Results from the linear regression analysis to test if the slope of the relationship between overall cortical activity and the relative contribution of corticocortical inputs depends on the experimental manipulation and tonotopic position. Significance codes: 0 ‘***’; 0.001 ‘**’; 0.01 ‘*’; 0.05 ‘.’

| | | $\beta$ Estimate | Std. Error | t value | Pr(>\|t\|) |
| --- | --- | --- | --- | --- | --- |
|  | (Intercept) | 4.6706 | 0.6006 | 7.777 | 1.23e-13 *** |
| Contrast to before trauma | Trauma | 1.3112 | 0.6923 | 1.894 | 0.05921 . |
|  | Recovery | -2.0402 | 0.7644 | -2.669 | 0.00803 ** |
| Contrast to below trauma region | Trauma | 1.5460 | 0.7996 | 1.934 | 0.05411 . |
|  | Above | 0.8681 | 0.6746 | 1.287 | 0.19915 |

**Table 2:** Linear regression of offset. results from the linear regression analysis to test if the offset of the relationship between overall cortical activity and the relative contribution of corticocortical inputs depends on the experimental manipulation and tonotopic position. Significance codes: 0 ‘***’; 0.001 ‘**’; 0.01 ‘*’; 0.05 ‘.

| | | $\beta$ Estimate | Std. Error | t value | Pr(>\|t\|) |
| --- | --- | --- | --- | --- | --- |
|  | (Intercept) | 0.55181 | 0.05748 | 9.601 | < 2e-16 *** |
| Contrast to before trauma | Trauma | -0.11058 | 0.06626 | -1.669 | 0.09617 . |
|  | Recovery | -0.06962 | 0.07316 | -0.952 | 0.34205 |
| Contrast to below trauma region | Trauma | 0.03474 | 0.07652 | 0.454 | 0.65012 |
|  | Above | 0.17333 | 0.06456 | 2.685 | 0.00767 ** |

### Response tuning revealed differences between acute and long-term tonotopic reorganization

We presented varying pure tone frequencies of moderate intensity (44 and 64 dB SPL) before and acutely after the sound trauma, and after 4-6 weeks of recovery. Thereby we characterized the frequency tuning of each individual recording position. In general, acutely after the trauma we found a prominent decrease of tuning width due to the vanished responses to the mid frequency range of 2 – <8 kHz in all recordings. Even at recording positions within the tonotopic representation of the mid frequency range no responses could be measured acutely after the sound trauma. After weeks, mid frequency responses only partially recovered. **Fig. 3A** shows a representative example with vanished responses after the trauma from 2 kHz (trauma frequency) upwards (including the BF) based on the RMS value of the AvgRecCSD (top). After weeks, tone-evoked RMS amplitudes of AvgRecCSD (top) and RelResCSD (bottom) largely recovered, even for the trauma frequency. Before trauma induction, BF sites were equally distributed across the frequency ranges below (<2 kHz), around (2 – <8 kHz), and above (≥8kHz) the trauma frequency for 44 dB SPL (*Pre*; Chi-square: 3.06, df: 2, p=0.216) and 64 dB SPL (Chi-square: 0.06, df: 2, p=0.969). After 4-6 weeks of recovery, we found significantly fewer sites with a BF in the range of 2-8 kHz with significant effects for stimulation with 64 dB SPL (*Recovery*; (1) 44 dB SPL: Chi-square: 0.115, df: 2, p=0.11; (2) 64 dB SPL: Chi-square: 8.33; df: 2, p=0.015; see **Fig. 3A** for an example). Only in 1 individual example, the BF in the recovery measurement was in the range around trauma (2 – 8 kHz), which we therefore did not consider for statistical analyses (**Fig. 3B**). For analyzing response bandwidths before (*Pre*) and after (*Trauma*) the trauma and 4-6 weeks later (*Recovery*), Q40dB values (in octaves) were calculated for the RMS value of the AvgRecCSD and RelResCSD. AvgRecCSD and RelResCSD response bandwidths showed significant reduction acutely after trauma induction in all 3 tonotopic site categories, viz. tonotopic regions with BFs below, near and above the trauma frequency. A full recovery of response bandwidths was found at recording sites below or above the trauma after 4-6 weeks of recovery, but not at the trauma site (**Fig. 3C**).

Based on the RMS values of AvgRecCSD and RelResCSD obtained after stimulation at moderate sound level (64 dB SPL) evoked responses were divided into sound frequency bins below, at and above the trauma frequency (**Fig. 3D;** note that we distinguish between *trauma site* which is the recording site at the tonotopic position of the trauma frequency (cf. the three columns of **Fig. 3D**), and *trauma frequency* which is the stimulation frequency used to induce the trauma (cf. abscissa values in the graphs in each column of **Fig. 3D**). After trauma induction, AvgRecCSD was significantly reduced for stimulation frequencies near and above the trauma frequency, but not below the trauma frequency, irrespective of the recording site. This is reflected by a 2-factorial rmANOVA for data obtained at each recording site that revealed main effects for the factor “stimulation frequency” and an interaction of factors “time point *X* stimulation frequency” for recording sites below the trauma, main effects for “frequency” at recording sites at the trauma, and mainly interaction effects at sites above the trauma (**Tab. 3.1-1.3**). Post-hoc Bonferroni-corrected Student’s t-tests were used to investigate statistical difference between paired samples. RelResCSD was most significantly reduced at the trauma site for frequencies near the trauma frequency (2-factorial rmANOVA with main effect of factor “stimulation frequency” (**Tab. 1.5**). After 4-6 weeks of recovery, AvgRecSD and RelResCSD were significantly increased for most stimulation frequencies in tonotopic regions below the trauma (main effects for the factor “stimulation frequency” and an interaction of factors “time point *X* stimulation frequency”; **Tab. 3.4**). No significant changes of both parameters were found before the trauma and after 4-6 weeks at BF sites higher than the trauma indicating a full recovery (no sig. effects for the rmANOVA tested on the RelResCSD at BF high).

**Table 3.** Report of significant repeated measures ANOVA effects for CSD parameters. rmANOVAs were Huyn-Feldt corrected and based on a significance level of α*=0.05

| FACTOR | F-value | p-value |
| --- | --- | --- |
| **1. Figure 3D; 2-way rmANOVA to test for AVREC RMS for "time point" and "frequency" at BF low (2-way rmANOVA; $\alpha^*=0.05$)** |  |  |
| "stimulation frequency" | $F_{2,24}=0.86$ | $p<0.001$ |
| "stimulation frequency x time point" | $F_{4,48}=4.62$ | $p=0.007$ |
| **2. Figure 3D; 2-way rmANOVA to test for AVREC RMS for "time point" and "frequency" at BF mid (2-way rmANOVA; $\alpha^*=0.05$)** |  |  |
| "stimulation frequency" | $F_{2,24}=25.59$ | $p<0.001$ |
| **3. Figure 3D; 2-way rmANOVA to test for AVREC RMS for "time point" and "frequency" at BF high (2-way rmANOVA; $\alpha^*=0.05$)** |  |  |
| "stimulation frequency x time point" | $F_{4,48}=2.83$ | $p=0.042$ |
| **4. Figure 3D; 2-way rmANOVA to test for RelRes RMS for "time point" and "frequency" at BF low (2-way rmANOVA; $\alpha^*=0.05$)** |  |  |
| "stimulation frequency" | $F_{2,24}=1.53$ | $p=0.016$ |
| "stimulation frequency x time point" | $F_{4,48}=3.28$ | $p=0.033$ |
| **5. Figure 3D; 2-way rmANOVA to test for RelRes RMS for "time point" and "frequency" at BF mid (2-way rmANOVA; $\alpha^*=0.05$)** |  |  |
| "stimulation frequency" | $F_{2,24}=21.91$ | $p<0.001$ |
| **6. Figure 3D; 3-way rmANOVA to test for layer-specific tuning shifts at BF low (3-way rmANOVA; $\alpha^*=0.05$)** |  |  |
| "Sink" | $F_{3,36}=14.1$ | $p<0.001$ |
| "stimulation Frequency" | $F_{2,24}=10.6$ | $p<0.001$ |
| "stimulation frequency x time point" | $F_{4,48}=5.38$ | $p=0.0051$ |
| "Sink x stimulation frequency" | $F_{6,72}=3.53$ | $p=0.042$ |
| "Sink x stimulation frequency x time point" | $F_{12,144}=2.87$ | $p=0.046$ |
| **7. Figure 3D; 3-way rmANOVA to test for layer-specific tuning shifts at BF mid (3-way rmANOVA; $\alpha^*=0.05$)** |  |  |
| "Time point" | $F_{2,24}=3.47$ | $p=0.047$ |
| "stimulation frequency" | $F_{2,24}=3.74$ | $p=0.039$ |
| "Sink x stimulation frequency x time point" | $F_{12,144}=4.15$ | $p=0.007$ |
| **8. Figure 3D; 3-way rmANOVA to test for layer-specific tuning shifts at BF high (3-way rmANOVA; $\alpha^*=0.05$)** |  |  |
| "Sink" | $F_{2,30}=55$ | $p<0.001$ |
| "stimulation frequency x time point" | $F_{2,30}=72.6$ | $p<0.001$ |

### Layer-specific changes of synaptic circuits underlying compensatory tonotopic shifts

In order to reveal layer-specific changes of synaptic processing in AI, CSD profiles at moderate sound level (64 dB SPL) were further analyzed directly before and after trauma induction as well as after 4-6 weeks.. **Fig. 4A** shows a representative example of effects of sound trauma induction on early sinks in granular layers III/IV (S1) and infragranular layers V (iS1), and subsequent sinks in supragranular layers I/II (S2) and infragranular layers VI (S3). Within subjects, CSD profiles showed comparable BFs during the *pre/trauma* and *recovery* recordings at regions above and below the trauma. For recovery measurement in animals with recordings at BF sites around the trauma in the pre-condition, we compared cortical activity after stimulation with the BF measured before trauma induction (see above). Sinks S1 and iS1 are neuronal observables of early thalamocortical synaptic input and immediate local corticocortical amplification, while S2 and S3 are related to supragrananular and infragranular corticocortical synaptic populations, respectively (Stoelzel et al., 2008; Happel et al., 2010; Schaefer et al., 2015). Below trauma (< 2kHz), the given example shows no significant change of the BF-evoked CSD profile after noise exposure, but a general increase of synaptic current flow across the entire cortical column after recovery. At tonotopic patches around the trauma site, we found a highly significant reduction of all early and late sink components after the noise trauma. After 4-6 weeks, cortical activation at these regions showed again a typical canonical feedforward CSD profile with initial input in granular (S1) and infragranular (iS1) layers and subsequent translaminar activations (S2 and S3). However, tonotopic tuning generally shifted its BF/CF tuning away from the initial BF range around the trauma frequency between 2 and 8 kHz (**Fig. 4B**). At tonotopic sites above the trauma (> 8kHz), trauma induction did not lead to obvious immediate changes while stimulating with the BF. After the recovery a significant increase was observed mainly in early and late infragranular activity. Increased supragranular activation showed a similar trend as at sites with BFs below the trauma.

In order to quantify these circuit effects, we have analyzed sink peak amplitudes (**Fig. 4**). Sink peak amplitudes at recording sites below the trauma showed no significant change after noise induction (compare with **Fig. 3D**) except for a decreased granular sink amplitude for stimulation within the trauma frequency range. After recovery, at tonotopic regions below the trauma all four sink components were increased when stimulating with the BF (cf. frequency bin ‘Below T.’; 3-factorial rmANOVA with significant effects for main factor “sink”, but not for “time point” or “stimulation frequency”; **Tab. 3.6**).. Therewith, the layer-specific sink activity adaptation is in accordance with the columnar AvgRecCSD response, which was also increased after recovery at tonotopic sites below the trauma for all stimulation frequencies (cf. **Fig. 3D**). The subsequent CSD profile analysis revealed that this was due to significantly increased activation across all cortical layers for low frequency stimulation, and to mainly increased granular activation for high frequency stimulation (**Fig. 4B**). At tonotopic sites around the trauma (2 – <8kHz) the sound trauma generally reduced all sink peak amplitudes irrespective of the stimulation frequency acutely and long-term. Depending on peak amplitudes before the trauma induction, this decrease was significant (3-factorial rmANOVA with main effect of factor “time point” and “stimulation frequency” and their interaction with the factor “sink”; **Tab. 3.7**). For recovery measurements no data could be obtained as no BFs between 2 and 8 kHz were observed (cf. above). At tonotopic sites with higher BFs (≥8kHz), acutely after the trauma the granular sink interestingly showed an enhanced peak amplitude for stimulation within the trauma frequency range, in contrast to sites below the trauma. At both sides apart from the trauma region, we consistently found increased early infragranular input (iS1), which may reflect mainly local and BF-specific thalamocortical input. After recovery, early granular input was still significantly increased for middle frequency stimulation, but not for frequencies of the actual high-frequency BF-range. In contrast to “below trauma” sites, we only observed a trend of increased supragranular activation, but a significant increase of late infragranular activation (3-factorial rmANOVA with main effects of factor “sink” and interaction of “stimulation frequency x time point”; **Tab. 3.8**).

## Discussion

### Immediate effects of sound trauma on circuit-level activity in ACx

A huge body of literature has described neurophysiological changes throughout the auditory system following noise trauma (for a review see Eggermont, 2017a). Nevertheless, the temporal evolution of plastic processes within ACx has rarely been addressed. In this report we have described acute and long-term effects of noise trauma on functional neuronal circuitry in ACx. We demonstrated that exposing Mongolian gerbils to continuous, intense 2 kHz pure tones over 75 min led to increases in cortical activation thresholds across the hearing range of gerbils which recovered to a large extent over the course of 4-6 weeks leaving residual threshold increases of about 10 dB at the trauma frequency and slightly above (**Fig. 1C**). Following such trauma, we found that acutely the overall activity within ACx decreased while the relative contribution of inter-columnar corticocortical inputs to this overall activity increased (**Fig. 5**, middle panel). These observations are in line with the study by Novák et al. (2016) who report increased activity of different types of inhibitory interneurons in layers II/III and IV of ACx immediately after trauma which in addition to the reduced thalamic input would explain the overall activity reduction after trauma. Furthermore, they describe an unmasking of excitatory inputs in layer II/III only which might explain the relative increase of inter-columnar contribution.

### Chronic effects of sound trauma on circuit-level activity in ACx

After recovery from trauma, the overall activity of the ACx increased again and actually reached levels higher than during pre-trauma conditions which has been discussed to be a physiological correlate of subjective tinnitus (Noreña and Eggermont, 2003; Eggermont and Roberts, 2004), while the relative contribution of corticocortical inputs decreased to levels even below (for frequency regions above trauma) or equal to pre-trauma conditions (**Figs. 2-3**). This finding leads to the conclusion that the high overall activity in ACx after recovery must result from a restored or even increased thalamic activation (**Fig. 5**, right panel) which is in line with the increased activity in the medial geniculate body (MGb) reported after noise exposure (Kalappa et al., 2014). It seems likely that increased thalamic activation of ACx is the result of mechanisms of compensatory, homeostatic plasticity (Mossop et al., 2000; Schaette and Kempter, 2006; Noreña, 2011; Tighilet et al., 2016) or stochastic resonance (Krauss et al., 2016, 2018) within the subcortical auditory pathway (Eggermont, 2017a). This conclusion is further supported by our layer-specific analysis, demonstrating that early sink activity in both thalamocortical-recipient layers III/IV and Vb/VIa was significantly increased after recovery (**Fig. 4B**). However, the increased gain was limited to local intracolumnar circuits and is not broadcast to wide-spread corticocortical circuits. This seems in line with earlier observations which demonstrated that higher activity after noise trauma was restricted to regions with changed tonotopy (Seki and Eggermont, 2003). Our results suggest an overcompensation of altered frequency input in auditory cortex that is limited to local circuits which might impact on crosscolumnar spectral integration across the tonotopic gradient (for a schematic summary of our findings refer to **Fig. 5**).

### Plasticity in local cortical circuits and their long term compensation

In order to reveal frequency tuning changes across ACx, we analyzed the trauma induced plasticity of tuning functions at different tonotopic regions. Our results have revealed stronger overall cortical activation several weeks after the trauma at tonotopic sites with BF representations lower than the trauma frequency (**Fig. 3D**, *top left*) which was accompanied by an increase of the relative residual CSD (**Fig. 3D**, *bottom left*). At tonotopic high-frequency regions, overall columnar activation was moderately increased irrespective of the stimulation frequency (**Fig. 3D**, *top right*), without an increase of relative corticocortical contributions (**Fig. 3D**, *bottom left*) in line with the aforementioned overcompensation (**Fig. 2**).

The quantitative analysis of the individual sink components further showed that this increase in spectral representation at tonotopic sites below the trauma was mediated mainly by synaptic inputs in granular layers (S1) and corticocortical inputs in supragranular layers (S2). Thalamocortical inputs in infragranular layers (iS1) were only increased for BF stimulation (**Fig. 4B**, *left column*). Such a significant increase of broad spectral input in granular and supragranular layers was absent at tonotopic regions above the trauma (**Fig. 4B**, *right column*). Nevertheless, in regions below the trauma early infragranular sink activity was significantly enhanced for BF-stimulation. This difference of early inputs in granular and infragranular layers might indicate a differential sound trauma sensitivity of the tuning width for thalamocortical inputs in granular layers and collaterals in deeper layers yielding different contributions to cortical tuning (Constantinople and Bruno, 2013). Interestingly, the observation of a reduction of perineuronal nets in mice AI after acoustic trauma (Nguyen et al., 2017) may be a prerequisite for all the chronic plastic changes described in our report as they presumably require adaptations in synaptic connectivity. Considering the finding of compensated (below trauma) or even overcompensated (above trauma) relative contributions of lateral input, the described long-term effects on cortical frequency tuning might underlie a mainly local intracortical gain of afferent synaptic activity compensating the acute increase of relative crosscolumnar activity.

In summary, this study has demonstrated that pure-tone-induced sound trauma does not only alter net cortical activation, but leads to a change in the relative contributions of thalamocortically relayed and intracortically relayed activity. The reduction of cortical response amplitudes to pure-tone stimulation seen directly after trauma induction, is accompanied by an acutely increased relative increase of intracortically relayed input. After several weeks of recovery, when thalamic input tuned to the trauma frequency is increased again, the relative contribution of intracortically relayed input is decreased, at least to levels found before trauma induction.

## Author contribution

MJ, MFKH, HS and FWO designed the research; MJ, MFKH performed all experiments; MJ, MFKH analyzed the data; KT provided new reagents/tools; MJ, MFKH, KT, PK, AS, HS and FWO discussed the data; MJ, MFKH and HS prepared figures and wrote the paper

## Acknowledgements

We thank Kathrin Ohl for technical assistance. This work was supported by grants from the Deutsche Forschungsgemeinschaft (DFG) to FWO. The funders had no role in study design, data collection and analysis, decision to publish, or preparation of the manuscript.

